# Annotations of four high-quality indigenous chicken genomes identify more than one thousand missing genes in sub-telomeric regions with high G/C contents

**DOI:** 10.1101/2024.01.08.574549

**Authors:** Siwen Wu, Tengfei Dou, Sisi Yuan, Shixiong Yan, Zhiqiang Xu, Yong Liu, Zonghui Jian, Jingying Zhao, Rouhan Zhao, Xiannian Zi, Dahai Gu, Lixian Liu, Qihua Li, Dong-Dong Wu, Zhengchang Su, Junjing Jia, Changrong Ge, Kun Wang

## Abstract

**Background:** Although multiple chicken genomes have been assembled and annotated, the number of protein-coding genes in chicken genomes is still uncertain due to the low quality of these genome assemblies and limited resources used in gene annotations.

**Results:** To fill the gap, we annotated our four recently assembled high-quality genomes of four indigenous chickens with distinct traits using a combination of RNA-seq- and homology-based approach. Our annotated genes in the four chickens recovered 51 of the 274 “missing” genes in birds in general and 36 of the 174 “missing” genes in chickens in particular. Intriguingly, based on deeply sequenced RNA-seq data collected in multiple tissues in each chicken breed, we found a total of 1,420 new protein-coding genes in the four chicken genomes, which were missed in the reference chicken genome annotations. These newly annotated genes (NAGs) tend to have high G/C contents and be located in sub-telomeric regions of almost all assembled chromosomes and some unplaced contigs. The NAGs showed tissue-specific expression and we were able to verify 39 (92.9%) of 42 randomly selected ones in various tissues of the four chicken breeds using RT-qPCR experiments. We found that most of the NAGs also are encoded in previously assembled chicken genomes. The NAGs form functional modules with homology-supported genes that are involved in many important biological pathways. We also identified numerous unique genes in each indigenous chicken genome that might be related to the unique traits of each breed.

**Conclusion:** The ubiquitous presence of the NAGs in various chicken genomes indicate that they might play critical roles in chicken physiology. Counting these new genes, chicken genomes harbor more genes than originally thought.

## Background

Chicken (*Gallus gallus*) provides us with most protein sources in our daily life and also is a model organism to study the development, immunity and diseases of vertebrates [1]. Although multiple versions of chicken genomes have been assembled, such as those for the red jungle fowl (GRCg1∼GRCg6a), the broiler (GRCg7b) and the layer (GRCg7w), understanding of chicken genetics, egg and meat production, domestication and evolution is still limited due to the incomplete assemblies and annotations of these genome versions as well as the unavailability of high-quality genome assemblies of diverse indigenous chickens and their annotations. For example, the number of protein-coding genes encoded in chicken genome is still an issue of debate. On one hand, like other birds, chickens have a small genome that is about a third of other tetrapods’ genomes in size, resulted from large scale segmental deletions in the avian lineage during evolution [2], leading to a large number of gene loss, thus chickens might have fewer genes than other tetrapods [3]. On the other hand, dozens to hundreds of genes that are essential in other tetrapods are believed missing in chickens and other birds’ genomes due to incomplete assemblies of chicken genomes, in particular, the micro-chromosomes where both gene density and G/C contents are higher. More recently, Li et al [4] assembled a chicken pan-genome based on genomic data from 20 diverse breeds and identified a total 1,335 new genes. However, more than half of these new genes are micro-open reading frames (ORFs) with a coding DNA sequence (CDS) shorter than 300 bp, casting doubts on the authenticity of these “new genes”.

We recently sequenced and assembled genomes of four indigenous chicken breeds with unique morphological traits from Yunnan province, China, including Daweishan, Hu, Piao and Wuding chicken, using a combination of long reads, short reads, and Hi-C reads [5]. These chromosome-level assemblies are of higher or comparable quality with the recently released chicken reference genomes GRCg7b/w, providing us an opportunity to survey the repertoire of genes, particularly, protein-coding genes encoded in genomes of diverse chicken breeds. Using a pipeline that combines homology-based and RNA-seq-based methods, we identified a total of 1,420 new protein-coding genes that were missed in the annotations of the reference genomes GRCg6a and GRCg7b/w but encoded in at least one of the four indigenous chicken genomes. The newly annotated genes (NAGs) are much longer than and overlap only a small portion of the so-called new genes identified by Li et al [4]. Most of the randomly selected NAGs can be verified by RT-qPCR experiment, thus these NAGs are likely authentic. Counting these NAGs, chickens have a similar number of protein-coding genes as other tetrapods do.

## Methods

### Materials

The GRCg6a, GRCg7b, GRCg7w and quail (*coturnix japonica*) genomes and annotation files were downloaded from the NCBI Genbank with accession numbers GCF_000002315.6, GCF_016699485.2, GCF_016700215.2 and GCF_001577835.2. Our previously assembled four indigenous chicken genomes were downloaded from the NCBI Genbank with the BioProject number PRJNA865263. All the Illumina short DNA sequencing reads, RNA-seq reads of different tissues of the four indigenous chickens were downloaded from the NCBI SRA database with accession number PRJNA865247. The sequences of 8,338 essential avian proteins were obtained from the BUSCO aves_odb10 database [26].

### Real-time quantitative PCR (RT-qPCR) analysis

RT-qPCR was performed using the Bio-Rad CFX96 real-time PCR platform (Bio-Rad Laboratories. lnc, America) and SYBR Green master mix (iQTM SYBRGreen ® Supermix, Dalian TaKaRa Biotechnology Co. Ltd. Add). The primers of the 42 randomly selected putative new genes are listed in Supplementary Note. The β-actin gene was used as a reference. Primers were commercially synthesized (Shanghai Shenggong Biochemistry Company P.R.C). Each PCR reaction was performed in 25 μl volumes containing 12.5 μl of iQ™ SYBR Green Supermix, 0.5 μl (10 mM) of each primer, and 1 μl of cDNA. Amplification and detection of products was performed with the following cycle profile: one cycle of 95 °C for 2 min, and 40 cycles of 95 °C for 15 s, annealing temperature for 30 s, and 72 °C for 30 s, followed by a final cycle of 72 °C for 10 min. The specificity of the amplification product was verified by electrophoresis on a 0.8% agarose gel and DNA sequencing. The 2^−ΔCt^ method was used to analyze mRNA abundance. All samples were analyzed with at least three replicates, and the mean of these measurements was used to calculate mRNA expression.

### Protein-coding gene annotation

To annotate the protein-coding genes in the assembled indigenous chicken genomes, we masked the repeats in each genome using WindowMasker (2.11.0) [27], and then we annotated the protein-coding genes using a combination of homology-based and RNA-based methods. For homology-based annotation, we collected all the protein-coding genes, pseudogenes and their corresponding CDS isoforms or exons in GRCg6a, GRCg7b and GRCg7w as the templates. The protein-coding genes in these three assemblies were recently predicted by the NCBI eukaryotic genome annotation pipeline that uses a combination of mRNA- and protein-based homology methods and ab initio methods. We mapped all the CDS isoforms of protein-coding genes and exons of pseudogenes in GRCg6a, GRCg7b and GRCg7w to each of the assembled indigenous chicken genomes using Splign (2.0.0) [28]. For each template gene whose CDSs could be mapped to an assembled genome, we concatenated all the mapped CDSs, and checked whether the resulting sequence forms an intact ORF (the length is an integer time of three and contain no stop codon in the middle). If yes, we called it an intact gene. If the CDSs of a template gene can be mapped to multiple loci in an assembled genome, we consider the locus with the highest mapping identity. If a concatenated sequence did not form an intact ORF, i.e., the length is not an integer time of three (ORF shift) or it contains a stop codon in the middle (nonsense mutation), we mapped the DNA short reads from the same individual to the CDSs of the gene using bowtie (2.4.1) [29] with no gaps and mismatches permitted. If the sequence could be completely covered by at least 10 short reads at each nucleotide position, we consider the pseudogenization is fully supported by the short reads, and called the sequence a pseudogene; otherwise, we consider the pseudogenization is not supported by the short reads, and called it a partially supported gene, because the pseudogenization might be artificially caused by errors of the long reads that could not be corrected by our assembly pipeline. For a few selenoprotein template genes where the “opal” stop-codon UGA encode selenocysteine, we manually checked the mapped loci in our assemblies, and annotate them accordingly.

For RNA-based annotation, we first mapped all of the RNA-seq reads from various tissues as well as the mixture of tissues of the four chicken breeds into the rRNA database SILVA_138 [30] and filtered out the mapped reads. We then mapped the unaligned reads to each of the four indigenous chicken genome assemblies using STAR (2.7.0c) [31]. Based on the mapping results, we assembled transcripts in each chicken using Trinity (2.8.5) [32] with its genome-guided option. Next, we mapped the assembled transcripts in each chicken to its assembled genomes using Splign (2.0.0) [28], and removed those that at least partially overlap non-coding RNA genes (see below), protein-coding genes or pseudogenes predicted by the homology-based method. For the remaining transcripts, if we could find a longest ORF with at least 300 pb, we called it a protein-coding gene. If multiple ORFs were found in a transcript, we selected the longest one.

Additionally, to detect protein-coding genes that might be not included in the four assemblies but expression in the tissues, we de novo assembled transcripts using Trinity (2.8.5) [32] with its de novo option using RNA-seq reads that could not be mapped to any of our four assembled genomes.

### RNA-coding gene annotation

We annotated tRNA, rRNA, miRNA, snoRNA, telomerase RNA and SRP RNA using infernal (1.1.2) [33] with Rfam (v.14) database [34] as the reference. In addition, we predicted an assembled transcript longer than 1,000 bp but lacking an ORF as a lncRNA.

### Function assignment for NAGs

We did the two-way hierarchical clustering of all genes (including NAGs) in the four chickens based on their expression levels in different tissues of each chicken using Euclid distance between expression vectors. For each NAG, we selected the smallest cluster that contains the NAG and at least one homology-supported gene with cluster distance smaller than 50. The gene function of the NAG assumed to be similar to those of the homology-supported genes that appear in the same cluster.

### Neighbor-joining tree construction

We mapped the 8,338 essential avian proteins from the BUSCO aves_odb10 database [26] to each of the seven chicken’s CDSs as well as the coturnix japonica’s CDSs using blastx (2.11.0) [35]. We selected the 6,744 genes with greater than 70% sequence identity with the essential avian proteins in each of the eight genomes to construct a neighbor-joining tree. Since it is hard to make multiple alignments for very long sequences, we evenly divided the genes in each bird into 68 groups (each contains about 100 genes) and concatenated CDSs in each group with fixed order. We then aligned the concatenated sequences of the same group in the eight birds using Clustal Omega (1.2.4) [36]. We finally concatenated the 68 multiple alignments and constructed a consensus neighbor-joining trees with 1,000 rounds of bootstrapping using Phylip (3.697) [37].

## Results

### More than one thousand protein-coding genes missed in reference annotations are found in four indigenous chicken genomes

By using a combination of homology-based and RNA-seq-based method, we predicted 17,497∼17,718 protein-coding genes in each of our recently assembled four indigenous chicken genomes [5] (Table 1). Interestingly, these numbers of annotated genes in the indigenous genomes are similar to those annotated in the RJF genome GRCg6a (17,485), but fewer than those annotated in the broiler GRCg7b (18,024) and the layer GRCg7w (18,016) genomes. Specifically, we predicted 16,917∼17,141 genes in each indigenous chicken genome based on homology to genes and pseudogenes annotated in GRCg6a, GRCg7b and GRCg7w (Materials and Methods). Of these homology-supported genes in each genome, 16,270∼16,668 have an intact ORF (intact genes) (Table 1), and 473∼647 contain either a nonsense or an ORF shift mutation that cannot be fully supported by short DNA reads from the chicken. It is highly likely that such “mutations” might be due to errors in long reads that cannot be corrected by the short DNA reads, we therefore refer these genes as partially supported genes (Table 1). The vast majority of intact genes (98.7%∼98.8%) and partially supported genes (97.9%∼99.2%) in each genome are transcribed in at least one of the tissues examined using RNA-seq (Tables S1∼S4), suggesting that at least most of the homology-supported genes are likely authentic. Interestingly, we predicted 6∼7 genes in each indigenous chicken genome based on homology to pseudogenes in GRCg6a and/or GRCg7b/w (Table 1). These genes have an intact ORF that is fully supported by short DNA reads, and 4∼7 are transcribed in at least one of the tissues examined (Tables S1∼S4), thus, they are likely to be functional. For example, the RJF pseudogene *LOC*107049240 at locus chr33:1757489∼1758423 with a point deletion is mapped to chr33:2240289∼2241224 of Piao chicken, encoding an intact ORF that is supported by 805 short DNA reads as well as large numbers of RNA-seq reads in multiple tissues (Figure 1a, Table S3). Notably, numbers of homology-supported genes in the indigenous chickens (16,917∼17,141) are smaller than those annotated in GRCg6a (17,485), GRCg7b (18,024) and GRCg7w (18,016), which is because 486∼622 genes annotated in GRCg6a and GRCg7b/w become pseudogenes in each of the four indigenous chickens (Table 1). Moreover, we predicted 83∼94 pseudogenes in the indigenous chickens based on homology to pseudogenes annotated in GRCg6a, GRCg7b and GRCg7w (Table 1).

**Figure 1.**
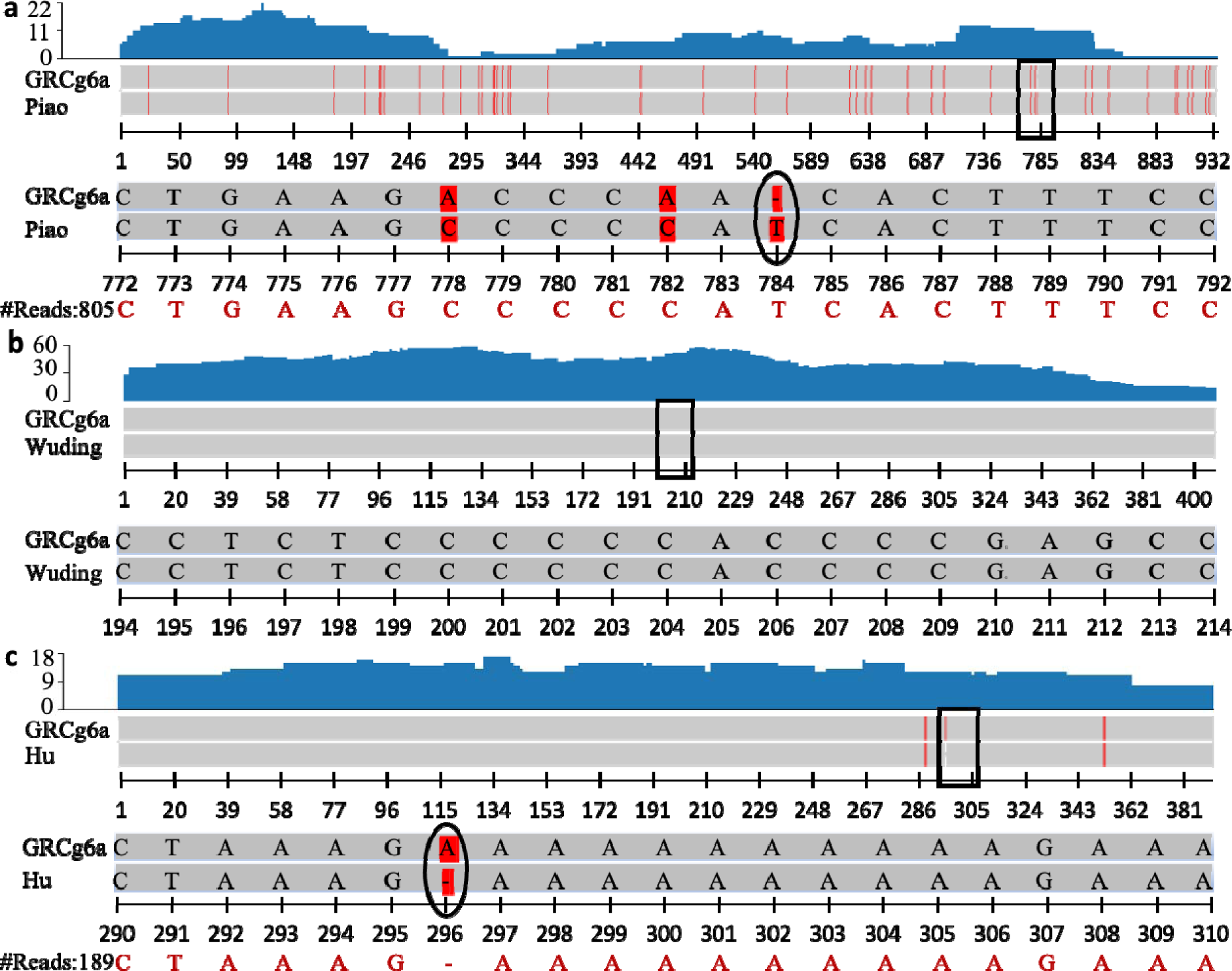
Examples of transcribed new genes and pseudogenes in the indigenous chickens and RJF. **a.** A protein-coding gene in Piao chicken is predicted based on a pseudogene LOC107049240 in RJF that harbors a point deletion of ‘T’ at position 784, leading to an ORF shift. **b.** A new gene predicted in Wuding chicken was also encoded in RJF. **c.** A new gene predicted in Hu chicken is pseudogenized in RJF, due to an insertion of ‘A’ at position 296, leading to an ORF shift.

**Table 1:**
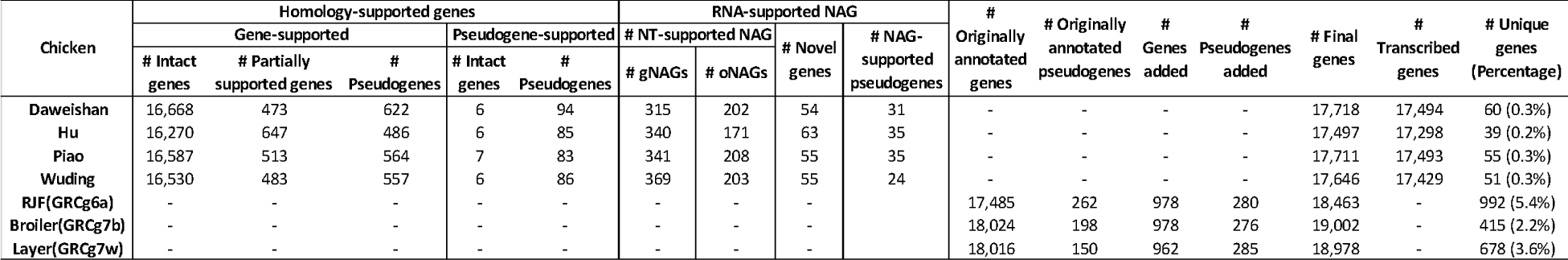
Summary of annotated protein-coding genes in the four indigenous chicken genomes in comparison with those in GRCg6a and GRCg7b/w.

Based on RNA-seq data that can be mapped to the four assembled indigenous chicken genomes, we identified 12,543∼13,930 putative genes in each of them (Table S5). Of these genes, 9,561∼10,596 at least partially overlap the homology-supported genes (Table S5), suggesting that homology-based method has already covered them. To avoid repeat, we do not consider them further. Of the remaining 2,982∼3,334 genes that do not overlap homology-supported genes, 571∼627 contain at least an ORF, and we consider them as NAGs (newly annotated genes) for further analysis (Table 1). Since some of these NAGs in the four assembled genomes are highly similar (identity > 98.5%) to one another, we removed the redundancy and ended up with a total of 1,420 unique NAGs in the four genomes, which were not annotated in GRCg6a, GRCg7b or GRCg7w. Interestingly, 24∼35 of the NAGs found in an indigenous chicken are pseudogenized in at least one of the other three chickens (Tables 1 and S6). Of the 1,420 NAGs, 1,277 (89.9%) (511∼572 in each of the four assembled genomes) are homologous to genes in the NT database (Table 1), so we refer them to as NT-supported NAGs. The remaining 143 (10.1%) (54∼63 in each assembled genome) NAGs do not have a known homolog in NT database, and thus, we consider them to be novel genes (Table 1). NT-supported NAGs in each breed (511∼572) are mapped to genes either in other breeds of *Gallus gallus* (315∼369 or 60.9%∼66.5%) rather than GRCg6a, GRCg7b and GRCg7w, or in other species (mainly avian species, 169∼208 or 33.1%∼39.7%) (Table S7), suggesting that they are likely true genes. Among the 1,277 NT-supported NAGs, 793 are mapped to genes in other chicken breeds, and we refer them to as *Gallus gallus*-supported NAGs (gNAGs). For the remaining 484 NT-supported NAGs that are mapped to species other than *Gallus gallus,* we refer them to as other species-supported NAGs (oNAGs). Combining the 484 oNAGs with the 143 novel genes, we identified a total of 627 new genes that have not been reported in chickens (Table S6).

### The NAGs show strong tissue-specific expression patterns and can be experimentally validated

As the first step to validate the NAGs, we examined their expression patterns using the RNA-seq data in ten tissues of each indigenous chicken (Materials and methods). As shown in Figures 2a∼2d and Tables S1∼S4, all the three groups of NAGs, i.e., gNAGs, oNAGs and novel genes, show strong tissue-specific expression patterns in all the four chickens, suggesting that they are likely authentic genes as we argued earlier. To further validate the 1,420 NAGs, we randomly selected 42 of them and quantified their transcription levels in various tissues of the four chicken breeds using RT-qPCR. Of the 42 selected NAGs, eight are novel genes, nine are oNAGs, and the remaining 25 are gNAGs. As shown in Tables S8∼S11, 21, 17, 10 and 15 of the 42 NAGs are encoded in Daweishan, Hu, Piao and Wuding chickens, of which 18 (85.71%), 14 (82.35%), eight (80%) and 15 (100%) were expressed in multiple tissues of the four breeds, respectively (Figures 3a∼3d). Combining the results from all the four breeds, 39 (92.86%) of the 42 selected NAGs were expressed in multiple tissues of at least one chicken breed, of which 23 (59.0%) are gNAGs, six (15.4%) are oNAGs and 10 (25.6%) are novel genes (Tables S8∼S11). Notably, the expression patterns in different tissues of the 39 new genes measured by RT-qPCR are not very similar to those quantified by RNA-seq reads (Figures 3a∼3d), which might be due to the different sensitivities of the two methods. Nonetheless, again, we conclude that most of our identified NAGs are likely authentic.

**Figure 2.**
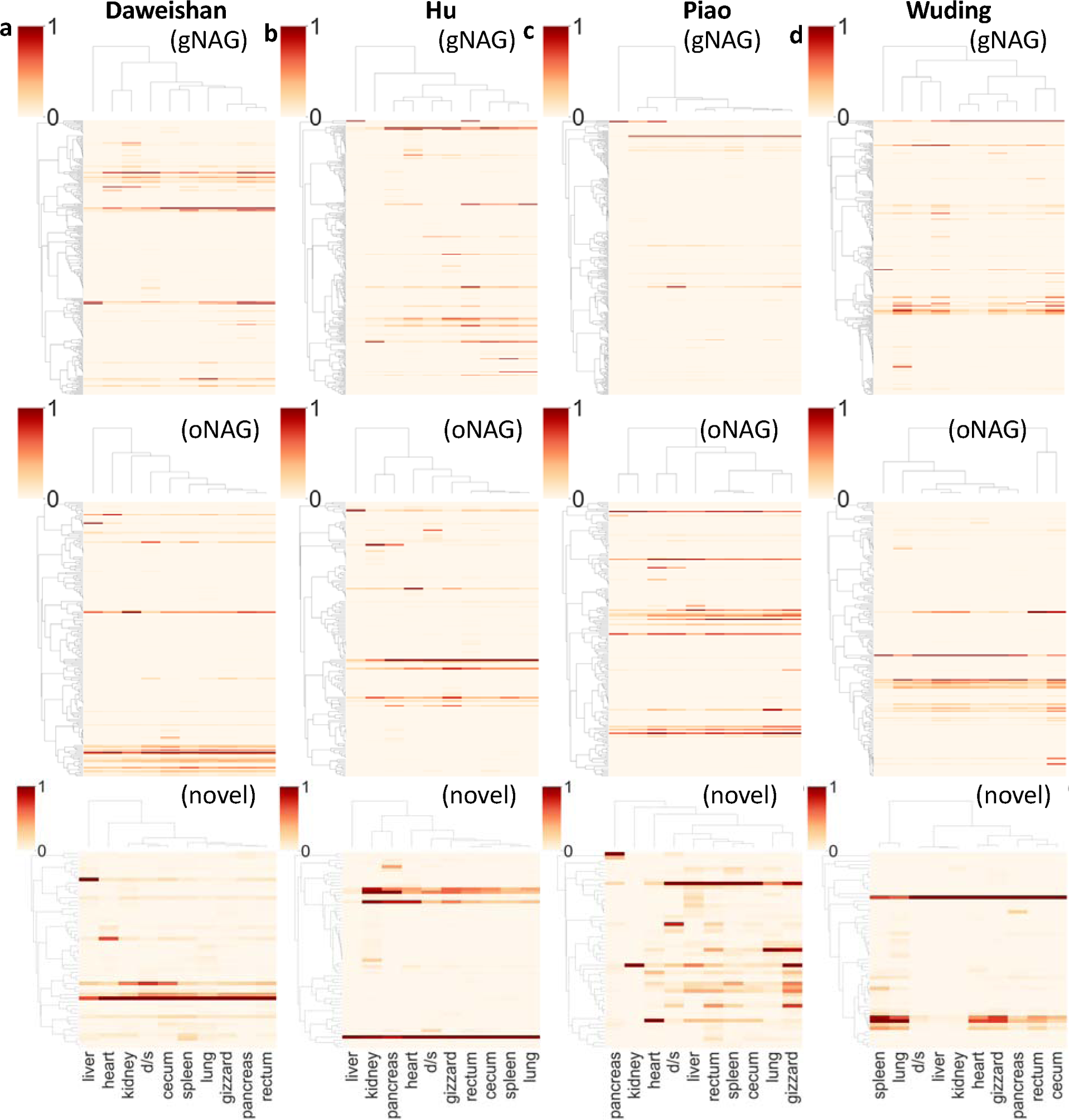
Expression levels of NAGs in different tissues of the four indigenous chickens from RNA-seq data. **a.** Expression levels of NAGs in different tissues of the Daweishan chicken. **b.** Expression levels of NAGs in different tissues of the Hu chicken. **c.** Expression levels of NAGs in different tissues of the Piao chicken. **d.** Expression levels of NAGs in different tissues of the Wuding chicken.

**Figure 3.**
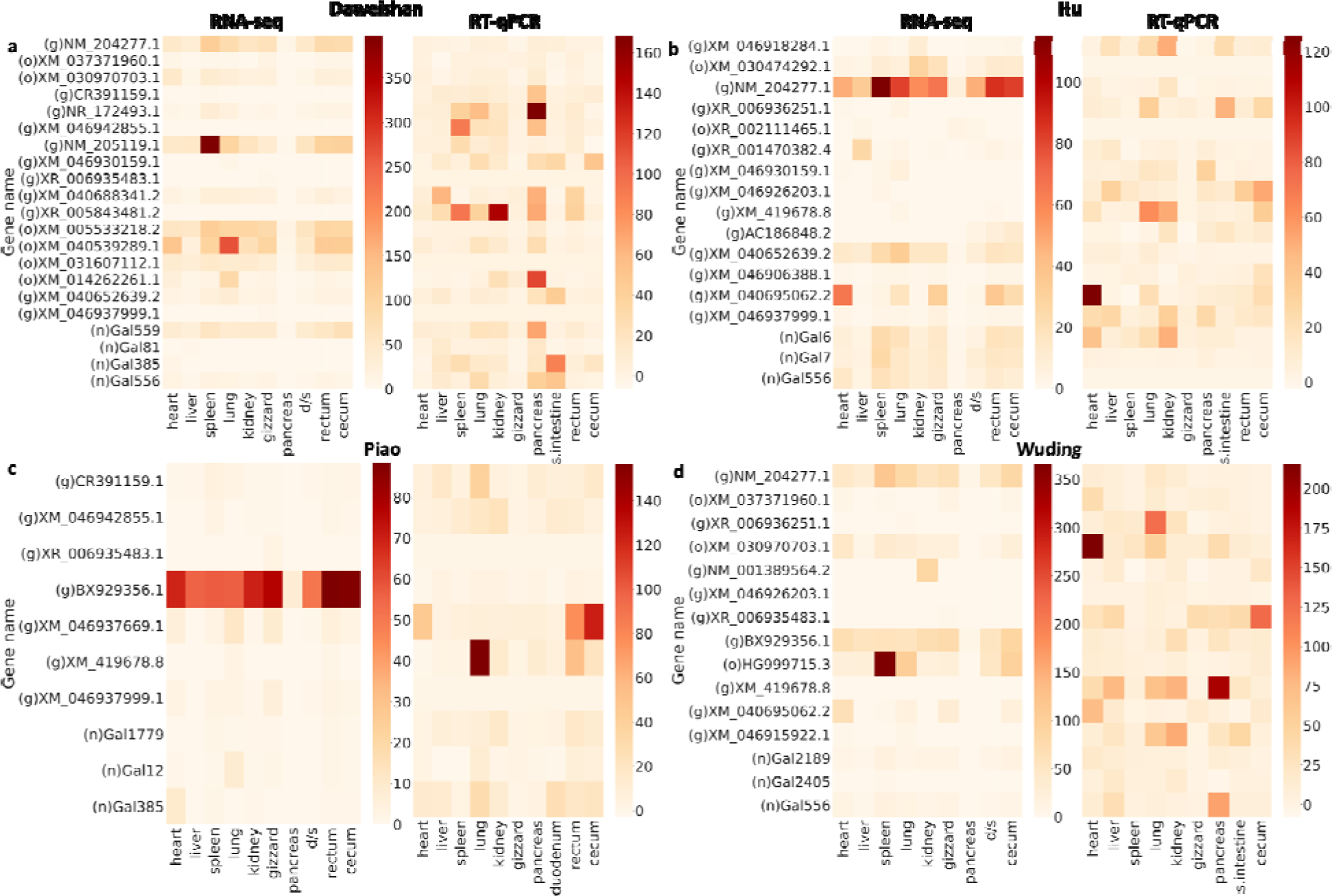
Heatmaps of expression levels of the 42 randomly selected NAGs in different tissues of the four indigenous chickens measured by RNA-seq and RT-qPCR. **a.** Expression levels of NAGs in different tissues of the Daweishan chicken. **b.** Expression levels of NAGs in different tissues of the Hu chicken. **c.** Expression levels of NAGs in different tissues of the Piao chicken. **d.** Expression levels of NAGs in different tissues of the Wuding chicken. In each subfigure, genes with ‘g’ labels represent gNAGs, genes with ‘o’ labels represent oNAGs, and genes with ‘n’ labels represent novel genes. The NAGs that are not encoded in a genome are not shown in the breed. “d/s” represents duodenum/small intestine.

### The NAGs have limited overlaps with the previously identified 1,335 new genes in chicken genomes

We compared the 1,420 NAGs with the 1,335 new genes recently found in 20 chicken genomes [4]. All the three groups of NAGs, i.e., gNAGs, oNAGs and novel genes, with a mean length of 8,232, 5,190 and 5,378 bp, respectively, are much longer than the 1,335 new genes with a mean length of only 1,413 bp (p=8e-137, 7e-85, 3e-22, respectively, Wilcoxon rank-sum test) (Figure 4a). One of the reason for the discrepancy is that 756 (56.6%) of the 1,335 new genes are mini-ORFs [6] with a CDS length of 100∼300 bp, while all of our 1,420 NAGs have a CDS length longer than 300 bp (Figure 4a), indicating that more than half of the earlier predicted so-called new genes might not be bona fide protein-coding genes. Of the 1,335 new genes, 660 (49.4%) can be mapped to at least one of the four indigenous chicken genomes with an identity greater than 98.5%. Of these 660 mapped so-called new genes, 246 overlap our 214 predicted genes in one of the four chickens, including 196 overlapping 176 homology-supported genes in one of the four chickens, and 50 overlapping 38 of our NAGs, while 246 (59.4%) of the remaining 414 so-called new genes are mini-ORFs, and thus were not predicted as genes by our pipeline. Of the remaining 675 (51.6%) of the 1,335 new genes, which cannot be mapped to any of the four indigenous genomes, 510 (75.6%) are mini-ORFs, while the other 165 (24.4%) are longer than 300 pb. Thus, though our 1,420 NAGs overlap 50 (3.7%) of the 1,335 so-called new genes, the two sets are quite different in terms of their lengths and overlapping rates.

**Figure 4.**
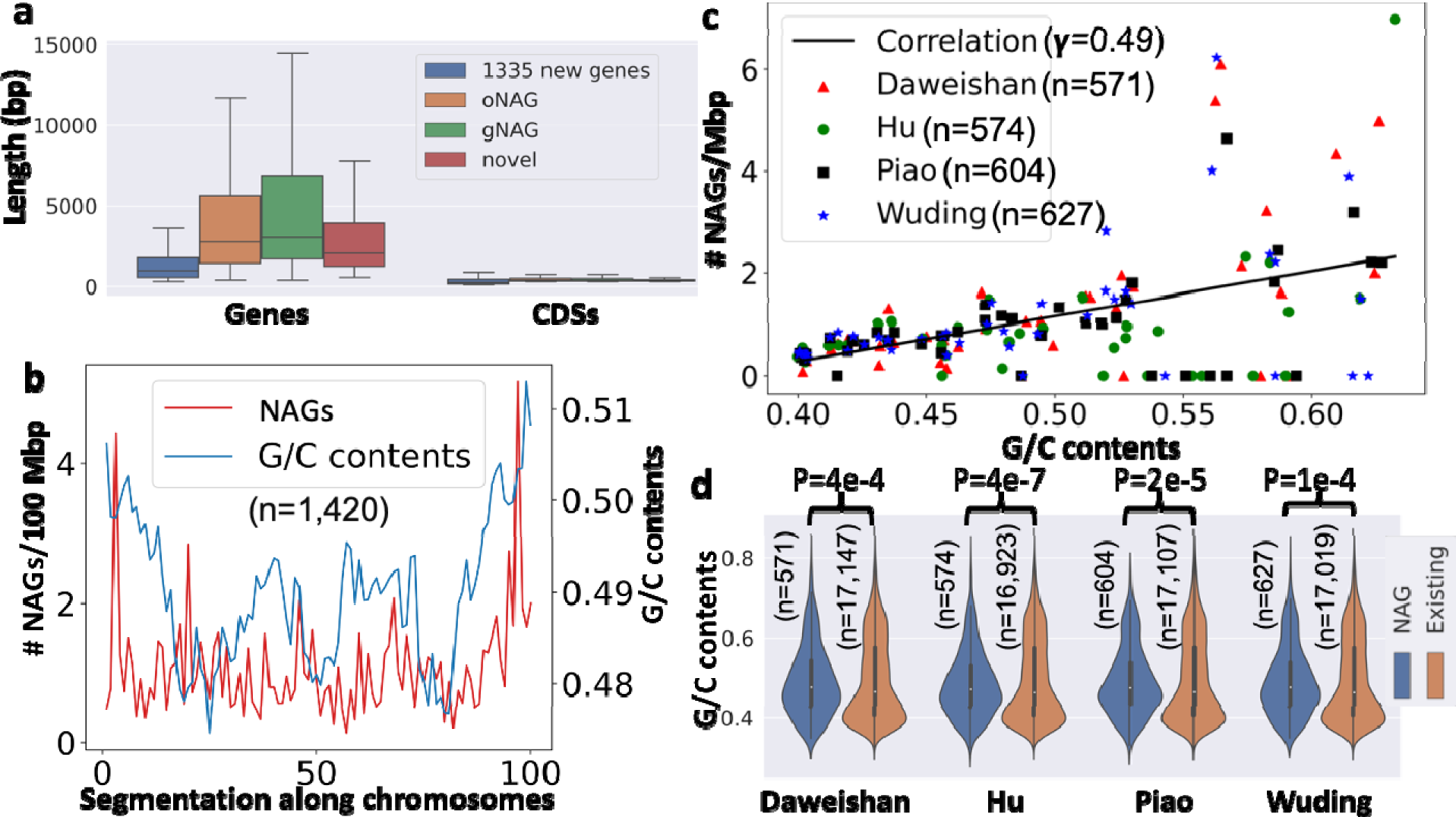
Some properties of our NAGs. **a.** Comparison of the lengths of our NAGs and their CDSs with those of the earlier predicted 1,335 new genes in 20 chickens. **b.** Average number of new genes per million bp and average G/C contents along evenly divided 100 segments of the 41 chromosomes in the four chicken genomes. **c.** The relationship between the number of new genes on a chromosome and its G/C contents in the four genomes. The black line is the linear regression of data from the four chickens. **d.** Comparison of G/C contents of the new genes with those of the existing genes.

### The NAGs are widely encoded in chicken genomes

To see whether the NAGs also exist in the previously assembled reference genomes but were simply missed by previous annotations, or they do not exist in these earlier assemblies, we mapped each of the 1,420 NAGs to the GRCg6a, GRCg7b and GRCg7w assemblies. To our surprise, 1,291 (90.9%) of the 1,420 NAGs could be mapped to at least one of the GRCg6a (1,258), GRCg7b (1,254) and GRCg7w (1,247) assemblies, including 760 gNAGs, 410 oNAGs and 121 novel genes. Of the 1,291 mapped NAGs, 1,142 have intact orthologous ORFs in at least one of the GRCg6a (978), GRCg7b (978) or GRCg7w (962) assemblies (Table 1). For examples, a NAG in Wuding chicken at chr11:17138466∼17139339 is mapped to RJF chr11:17570676∼17571083 that encodes an intact ORF (Figure 1b). Thus, most (1142, or 80.4%) of the 1,420 NAGs are encoded in these earlier assemblies but were missed by previous annotations (Table S6), due probably to the limited RNA-seq data used in annotation pipelines. Adding these intact NAG orthologs to the previously annotated lists of protein-coding genes, we increase the number of protein-coding genes in GRCg6a (18,463), GRCg7b (19,002) and GRCg7w (18,978) by 5.6% (978), 5.4% (978) and 5.3% (962), respectively (Table 1). Therefore, GRCg6a and GRCg7b/w encode more protein-coding genes than previously thought, supporting the previous conclusion inferred for birds in general [7]. On the other hand, 459 of the 1,291 mapped NAGs are pseudogenized in at least one of the GRCg6a (280), GRCg7b (276) and GRCg7w (285) assemblies (Table S6), i.e., they contain at least one nonsense or ORF shift mutations of normal genes. For example, a NAG in Hu chicken at locus chr2:128256077∼128257512 is mapped to RJF locus chr2:129144681∼129145071 with an insertion of ‘A’ supported by 189 short reads, leading to an ORF shift (Figure 1c). Thus, we also substantially increase the number of pseudogenes in GRCg6a (from 262 to 542), GRCg7b (from 198 to 474) and GRCg7w (from 150 to 435) (Table 1).

### The NAGs tend to be located at the sub-telomeric regions of the chromosomes with higher G/C contents

We examined the distributions of the NAGs along the chromosomes. As shown in Figures 5a∼5c, the NAGs are distributed on almost all the chromosomes, but have higher densities on micro-chromosomes and unplaced contigs (Figures 5d∼5f). The NAGs also tend to be located at the sub-telomeric regions of the chromosomes where G/C contents are higher (Figure 4b), consistent with a recent report [4]. This promoted us to analyze the relationship between the G/C contents in one Mbp genome region and the number of NAGs found in it. As shown in Figure 4c, the higher G/C contents of a genome region, the higher number of NAGs are found in it (Pearson correlation coefficient γ=0.49). Consistent with these facts and the higher G/C contents in micro-chromosomes, the NAGs tend to have higher G/C contents than the previously annotated genes (p<4e-4, K-S test) (Figure 4d).

**Figure 5.**
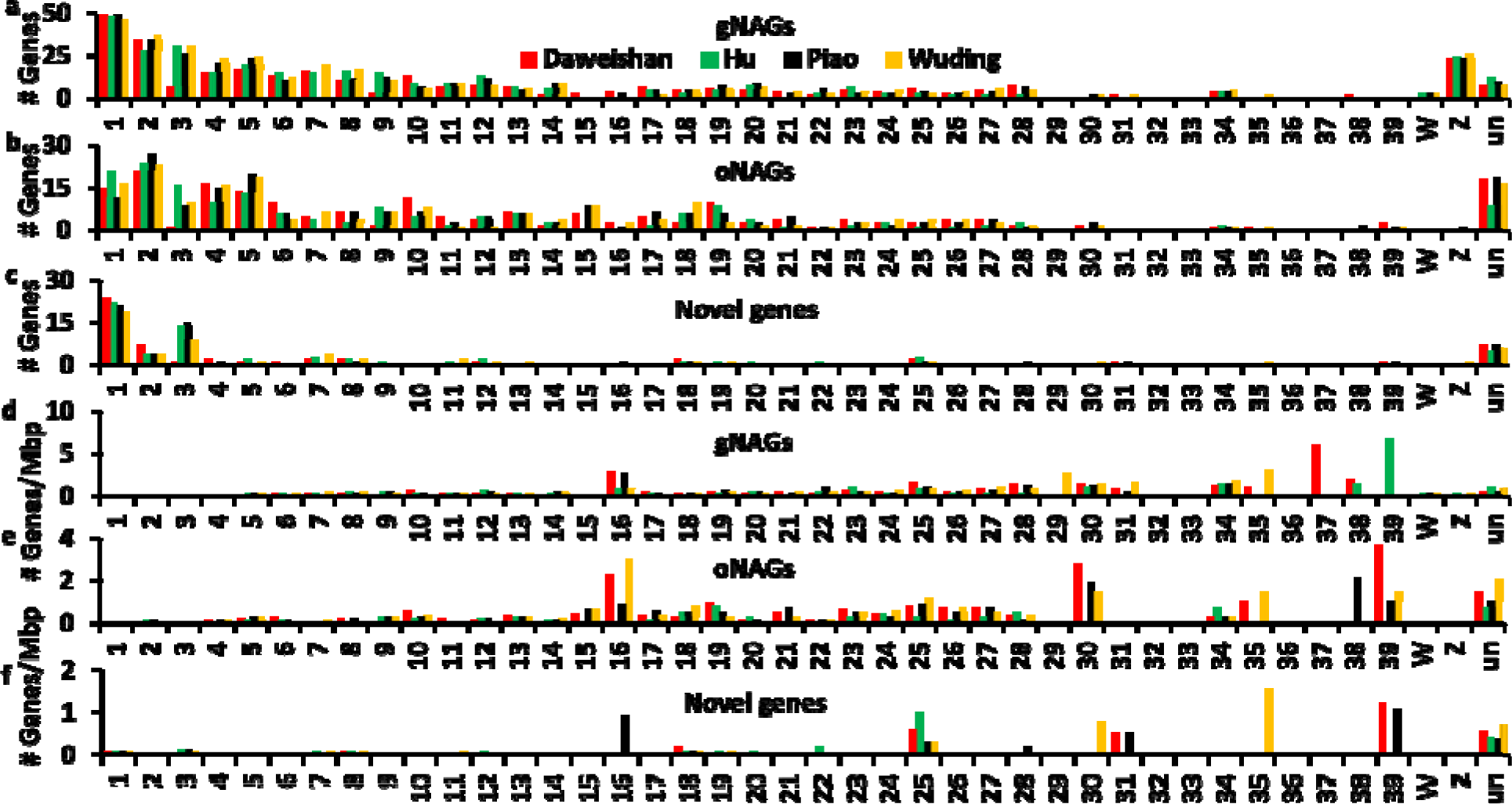
Distribution of different types of NAGs found in the four indigenous chicken genomes. **a∼c.** Number of gNAGs, oNAGs and novel genes found on each chromosome and unplaced contigs. **d∼f.** Number of gNAGs, oNAGs and novel genes per million bp found on each chromosome and unplaced contigs.

### The NAGs are involved in house-keeping functions

After demonstrating the wide distribution of the NAGs in chicken genomes, we asked whether there are any patterns for the NAGs to occur in these genomes. To this end, we clustered the chickens based on the similarity of presence, absence or pseudogenization of the 1,420 NAGs in the genomes. As shown in Figure 6a, two distinct cluster can be seen, one is formed by the four indigenous chickens and the other is formed by GRCg6a, GRCg7b and GRCg7w. Specifically, the former cluster is featured by the fact that only 40.2∼44.2% (571∼627) of the NAGs occur while the remaining 55.8∼59.8% (793∼849) are either absent or pseudogenized in the four indigenous chickens (Figure 6a, Table S6). In contrast, the latter cluster is featured by the fact that two thirds (962∼978) of the NAGs appear while the remaining one third (422∼458) are either absent or pseudogenized in GRCg6a, GRCg7b and GRCg7w. Moreover, we also clustered the NAGs based on their occurring patterns in the genomes and found that they are clustered in multiple distinct groups (Figure 6a). Although most of the NAGs do not have gene ontology (GO) [8] term assignments, those that have are involved in a total of 40 GO biological pathways (Table S12), many of which correspond to gene clusters shown in Figure 6a. These GO biological pathways are involved in house-keeping functions, including transcription regulation, signal transactions, immunity, cell growth, metabolism and apoptosis, to name a few (Table S12), indicating that many of them might be critical for chicken biology.

**Figure 6.**
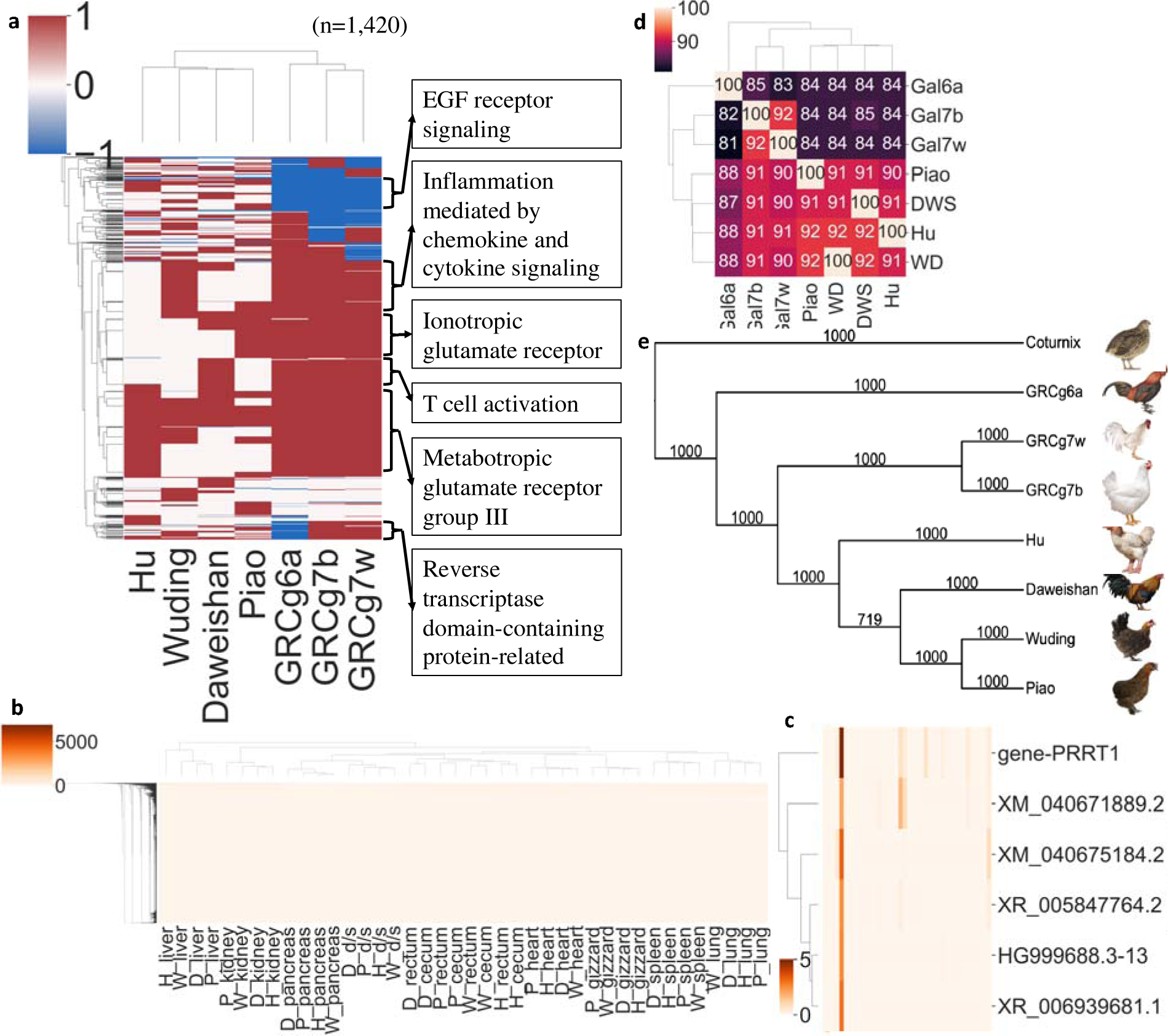
Occurring patterns of the 1,420 NAGs in the seven chicken genomes and phylogenetic relationships of the seven chickens. **a.** Heatmap of two-way hierarchical clustering of the new genes based on their appearance (1, brown), absence (0, white) and pseudogenization (-1, blue) patterns in the seven chicken genomes. **b.** Heatmap of two-way hierarchical clustering of expression levels of all genes in different tissues of the four chickens. **c.** An example of a cluster formed by five NAGs and gene *PRRT*1. **d.** Heatmap of two-way hierarchical clustering of the percentages of the genes that each chicken (rows) share with the others (columns). **e.** Consensus neighbor-joining tree, constructed using 6,744 essential protein-coding genes of the seven chickens and *coturnix japonica*.

### Most NAGs form tight clusters with homology-supported genes

As we mentioned above, most NAGs do not have GO term assignments. To reveal the possible functions for these NAGs, we performed a two-way hierarchical clustering on the expression matrix of the NAGs and all homology-supported genes in the four indigenous chickens in different tissues of each chicken. As shown in Figure 6b, the same types of tissues in the four chicken breeds are clustered together, indicating that the same types of tissues in different chicken breeds have similar transcriptome as expected. The genes also form numerous distinct clusters (Figure 6b). To infer the function of a NAG, we selected the smallest cluster containing the target NAG and at least one homology-supported genes with the Euclid distance from the leaves to the root of the cluster smaller than 50. We identified 1,118 such clusters containing 7,295 genes for 1,344 NAGs (Table S13), suggesting that most (94.6%) NAGs form a tight cluster containing at least one NAG and one homolog-supported gene. By “guilty by association”, each cluster might form a functional module. Thus, the NAGs might have similar functions as the homology-supported genes in the cluster. Figure 6c shows an example for a cluster formed by six genes including five NAGs and a homology-supported gene *PRRT1*. It is known that PRRT1 regulates basal phosphorylation level of glutamate receptor GRIA1, and promotes GRIA1 and GRIA2 translocation on cell surface [9, 10]. Thus, these five NAGs might also be involved in regulating GRIA1 and GRIA2 functions. The gene compositions of clusters for the other NAGs are shown in Table S13.

### Unique genes in each chicken breed might be related to its unique traits

Our more complete annotations of protein-coding genes in chicken genomes promoted us to analyze possible relationship between the gene contents and unique traits of the chickens. Notably, the broiler (GRCg7b) with 19,002 genes and the layer (GRCg7w) with 18,978 genes encode 1,260∼1,505 more genes than the four indigenous chickens with 17,497∼17,718 genes (Table 1). The broiler and the layer also encode 539 and 515 more genes than the RJF (GRCg6a) with 18,463 genes (Table 1). Some of the additional genes in the broiler and the layer genomes might be related to their meat and egg productions, respectively. Moreover, the RJF also encodes 745∼966 more genes than the four indigenous chickens, suggesting that the indigenous chickens might lose these genes during the domestication process. Furthermore, the four indigenous chickens share more genes with each other and with GRCg7b and GRCg7w than with GRCg6a, while GRCg7b and GRCg7w share more genes with each other than with GRCg6a and the four indigenous chickens (Figures 6d). These results suggest that the indigenous chickens are evolutionarily closer to one another and with GRCg7b and GRCg7w than with GRCg6a, and that GRCg7b and GRCg7w are more evolutionarily closer to each other than to GRCg6a. To confirm this argument, we constructed a neighbor-joining tree using 6,744 essential protein-coding genes (Table S14) in the seven chickens plus quail (*Coturnix japonica*) as the root. As shown in Figure 6e, the four indigenous chickens form a clade, the broiler (GRCg7b) and the layer (GRCg7w) form another clade, and the RJF (GRCg6a) branched earlier, confirming our conclusion.

We further analyzed the unique genes in each chicken genome. As shown in Tables 1 and S15), there are a varying number of unique genes in Daweishan (60), Hu (39), Piao (55), Wuding (51), GRCg6a (992), GRCg7b (415) and GRCg7w (678). Thus, the RJF has a far greater number of unique genes than the domesticated breeds, while the indigenous chickens have far smaller number of unique genes than the broiler and the layer. The unique genes in the indigenous chickens include varying number of NAGs, i.e., 19 gNAGs, 33 oNAGs and five novel genes in Daweishan chicken; 23 gNAGs, 12 oNAGs and three novel genes in Hu chicken; 20 gNAGs, 28 oNAGs and five novel genes in Piao chicken; 24 gNAGs, 20 oNAGs and six novel genes in Piao chicken. Although most unique genes in each chicken particularly those in the indigenous chicken do not have GO term assignments, those that do are involved in critical biological pathways (Table S16). For example, unique genes in Daweishan chicken with a miniature body size are involved in multiple pathways that might be related to its unique traits, including heterotrimeric G-protein signaling pathway-Gi alpha and Gs alpha mediated pathway, T cell activation, Ionotropic glutamate receptor pathway, Heterotrimeric G-protein signaling pathway-rod outer segment phototransduction, B cell activation and Metabotropic glutamate receptor group III pathway. Unique genes in Hu chicken with a very large body weight (∼6kg) are involved in FGF (Fibroblast growth factors) signaling, EGF (epidermal growth factor) receptor signaling and PDGF (platelet-derived growth factor) signaling pathways that play important roles in stem cell proliferation and growth. Unique genes in the layer (GRCg7w) that needs high calcium storage for the formation of eggshells are involved in sarcoplasmic/endoplasmic reticulum calcium ATPase 1 pathway for calcium assimilation. Unique genes in the broiler (GRCg7b) with fast growth rate are involved in multiple p53-related pathways, Wnt signaling pathway and glucose deprivation pathway. The RJF (GRCg6a) has the largest number (992) of unique genes that are involved in numerous pathways that might favor their wildlife, such as Wnt signaling pathway, B cell activation, de novo purine biosynthesis, TGF-beta signaling pathway, FAS signaling pathway, alpha adrenergic receptor signaling pathway and oxidative stress response. Besides the unique genes in each chicken breed that can be assigned GO term pathways, most of the unique genes can also be assigned functions based on the clusters that they are located in (Figure 6b, Table S13). For example, unique gene *Gal*574 (novel) in Daweishan chicken is in the same cluster with gene *CPSF1*, which might be related to the eye morphogenesis and the development of retinal ganglion cell projections to the midbrain [11]. Unique gene *XM_031614658.1* (oNAG) in Wuding chicken is in the same cluster with gene *MARF1*, which is a regulator of oogenesis required for female meiotic progression to repress transposable elements and preventing their mobilization, thus is essential for the germline integrity [12]. The unique genes in each chicken breed might be good candidates for experimental studies for their roles in forming the breed’s unique traits.

### “Missing” protein-coding genes are found in chicken genomes

It has been reported that 274 genes that are widely encoded in reptiles and mammals are missing in avian species in general and another 174 genes are missing in chicken (based on the galGal4 assembly) in particular [3]. However, some of these two sets of presumed missing genes have been recovered in chicken and other bird genomes [13–15]. We noted that 56 of the 274 missing genes in avian species and 43 of the 174 missing genes in chicken are annotated in at least one of the GRCg6a, GRCg7b or GRCg7w assemblies. To see whether our annotation recover any of the presumed missing genes, we mapped their human CDSs to each of the indigenous chicken genomes. We found that 48 and 36 of them, respectively, were among our predicted genes (Table S17). However, eight (ITPKC, TEP1, BRSK1, PLXNB3, MAPK3, HIGD1C, KMT5C and PACS1) of the 274 missing genes in avian species and seven (PPP1R12C, CCDC120, ATF6B, ASPDH, DDIT3, ADCK5, GPAA1) of the 174 missing genes in chicken, which were annotated in GRCg6a, GRCg7b or GRCg7w, did not appear in any of the four indigenous genomes (Table S17). It has been shown that RNA-seq data from multiple tissues can be used to recover presumed missing genes in birds [13–15]. To see whether more missing genes could be recovered by RNA-seq data collected in various tissues of the indigenous chickens, we *de novo* assembled transcripts using RNA-seq reads that could not be mapped to any of the four chicken genomes. Three of 34 such predicted CDSs/genes (Table S18) in the four chickens, can be mapped to three (MAPK3, SLC25A23 and HSPB6) of 274 missing genes in birds (Table S17A). MAPK3 is annotated in GRCg7b/w but missing in GRCg6a, while SLC25A23 and HSPB6 are missing in all these three previous annotations. Thus, loci of genes MAPK3, SLC25A23 and HSPB6 might be missed by the assemblies of the four indigenous chickens due probably to their refractory to the sequencing technologies used to assemble the genomes, and thus these genes are missed by our annotation pipeline.

### RNA genes in the chicken genomes

We also annotated RNA genes in each of the four indigenous chicken genomes. As summarized in Table S19, the four indigenous chicken genomes encode similar number of all the eight categories of RNA genes as annotated in GRCg6a and GRCg7b/w, except for snRNA, SRP (signal recognition particle) RNA and lncRNA. For snRNA, the four indigenous chicken genomes encode only three genes, while 66-75 were annotated in the GRCg6a and GRCg7b/w. For SRP RNA, the four indigenous chicken genomes encode 18∼19 genes, while only one was annotated in GRCg6a and GRCg7b/w. For lncRNA, the four indigenous chicken genomes encode 13,429∼15,080 genes, while only 8,233∼10,062 were annotated in GRCg6a and GRCg7b/w.

### Mitochondrial genomes of the chickens

We also annotated genes in our previously assembled mitochondrial genomes of the four indigenous chickens. We found 13 protein-coding genes on the mitochondrial genomes of Hu, Piao and Wuding chickens, which are the same as annotated in GRCg6a and GRCg7b (as a paternal genome, GRCg7w does not contain the mitochondrion). Interestingly, in the Daweishan mitochondrial genome, we found that ND5 (CDSs length=1,818bp in RJF) is a pseudogene because of an ORF-shift mutation caused by a five-bp deletion at CDS positions 1,552∼1,556, affecting 89 amino acids at the carboxyl terminus of the protein.

## Discussion

Although multiple improved versions of the RJF genome (GRCg1∼6a) [2, 16, 17] and commercial chicken genomes (GRCg7b/w) [18] have been assembled, understanding of chicken genomes is still far from complete. In particular, the number of protein-coding genes annotated in genomes of different breeds varies widely, and many presumed missing genes in chickens are still not found [3]. To better understand the gene repertoire in various chicken breeds and the underlying reasons of missing genes, in this study, by using a combination of homology-based and RNA-seq-based methods, we annotated the protein-coding genes in four indigenous chicken genomes assembled at chromosome-level with high-quality. We annotated similar number (17,494∼17,718) of genes in each of the indigenous chicken genomes as those previously annotated in the GRCg6a (17,485), GRCg7b (18,027) and GRCg7w (18,016). However, our RNA-seq-based annotation allowed us to uncovered from 511 to 572 new genes in each of the four indigenous chickens that are not seen in the reference annotations of GRCg6a and GRCg7b/w. After removing the repeated discoveries in different indigenous chicken genomes, we ended up with 1,420 NAGs (newly annotated genes), of which 143 might be novel genes with no homologs in the NT database, 484 are homologs to genes in other specie (most of which are avian) (oNAGs), and 793 are mapped to known chicken genes in other chicken breeds other than GRCg6a and GRCg7b/w (gNAGs). The tissue-specific expression of these NAGs and RT-qPCR validations of randomly selected NAGs in multiple tissues of at least one breed suggest that they are likely authentic. Combing the oNAGs and novel genes, we identified 627 new chicken genes. We compared our 1,420 NAGs with the 1,335 new genes recently reported by Li et al in chickens [4], and found that the two sets had only limited overlaps and differed largely in their lengths. The discrepancy might be due to different numbers of chickens/breeds used and different definitions of a gene adopted in the two studies. For example, more than half (756, 56.6%) of the 1,335 previously identified genes have a CDS length of only 100∼300 bp, thus they might be mini-ORFs [6] and bona fide protein-coding genes.

Moreover, we identified 48 of the 274 missing genes in birds in general and 36 of another 174 missing genes in chicken in particular in at least one of the four assembled indigenous genomes (Table S17). We also uncovered three of the 274 missing genes in birds (Table S17A) by mining unmapped RNA-seq reads. Two of the three recovered genes are missing in the GRCg6a and GRCg7b/w annotations. Our ability to identify the 1,420 NAGs and to recover missing genes indicates that the approach that we used to annotate the indigenous chicken genomes have largely overcome the limitations of the earlier methods. However, the assembled chr16 and some other micro-chromosomes of the four indigenous chickens are still not complete enough, thus, none of our 1,420 NAGs recall any of the 274 missing genes in birds in general and additional 174 missing genes in chickens in particular. Therefore, if the remaining presumed missing genes are encoded in the chicken genomes, accurate-enough ultra-long-reads (>100 kbp) from new sequencing technologies that allows the recent telomere to telomere assembly of a hypoploid human genome [19] might be needed to completely assemble the chicken genomes, thereby recovering the still missing genes.

Interestingly, 1,142 (80.4%) of the 1,420 NAGs also are encoded in at least one the GRCg6a, GRCg7b or GRCg7w assemblies (Table S6). The wide presence of the NAGs in these diverse chicken genomes indicate that most of the NAGs might play crucial roles in house-keeping functions as also indicated by GO pathways that they are involved in (Table S12) and gene functions that are assigned to them (Table S13). Including the 1,420 NAGs and recovered missing genes, we annotate 17,497∼17,718 protein-coding genes in the four indigenous chicken genomes and increase the number of protein-coding genes in GRCg6a, GRCg7b and GRCg7w to 18,463, 19,002, 18,978, respectively. Considering many gene-rich micro-chromosomes are still not fully assembled, the number of annotated genes may increase further once all the micro-chromosomes are fully assembled. Thus, the chicken genomes might encode similar number of protein-coding genes as other tetrapods (18,000∼25,000) [18, 20] such as humans[21], as previously suggested for birds in general [7]. The much larger number of protein-coding genes in the two commercial chickens (GRCg7b/w) than in RJF (GRCg6a) and indigenous chickens might be the results of the breeding programs used to produce the fast-growth and high egg-laying terminal commercial lines for production by taking advantage of hybrid vigor through a series of gene introgression from multiple purebred populations generated by intensive artificial selection [22–25].

## Conclusions

In this study, we annotated assembled high-quality genomes of four indigenous chickens using a combination of homology and RNA-seq based approach. We not only recovered dozens of previously presumed “missing” genes in chickens, but also found a total of 1,420 new genes that were missed by previous annotations. These new genes are often located in sub-telomeric regions and some unplaced contigs where G/C contents are high. We verified 39 (92.9%) of 42 randomly selected new genes using RT-qPCR experiments. The new genes showed tissue specific expression patterns and form functional modules with genes with known functions, and are involved in many important biological pathways. Most of the new genes also are encoded in the previous chicken genome assemblies. Counting these new genes, chicken genomes encode more genes than originally thought. We also identified numerous unique genes in each indigenous chicken genome that might be related to the unique traits of each breed.

## Declarations

### Ethics approval and consent to participate

All the experimental procedures were approved by the Animal Care and Use Committee of the Yunnan Agricultural University (approval ID: YAU202103047). The care and use of animals fully complied with local animal welfare laws, guidelines, and policies.

### Consent for publication

Not applicable.

### Availability of data and materials

The annotation code and pipeline description are available at https://github.com/zhengchangsulab/A-genome-assebmly-and-annotation-pipeline

### Authors’ contributions

CG, JJ and ZS supervised and conceived the project; KW, TD, SY^2^, ZX, YL, ZJ, JZ, RZ, XZ, DG, LL, QL and DW collected tissue samples and conducted molecular biology experiments; SW and SY^1^ assembled and corrected the genomes; SW and ZS performed data analysis; and SW, ZS and JJ wrote the manuscript.

## Acknowledgments

This work was supported by the Yunnan Agricultural University and Department of Bioinformatics and Genomics of the University of North Carolina at Charlotte.

