## Supplementary Note for "Annotations of four high-quality indigenous chicken genomes identify more than one thousand missing genes in sub-telomeric regions with high G/C contents"

**1 Gene name: XM_046918284.1 (NT-support)**

|  | Sequence (5'->3') | Tem. strand | Len(bp) | Start | End | Tm | GC% | Self com. | Self 3' com. |
| --- | --- | --- | --- | --- | --- | --- | --- | --- | --- |
| Forward primer | ACAAACTGAGAAAGGGGCGT | Plus | 20 | 4174 | 4193 | 59.82 | 50.00 | 3.00 | 0.00 |
| Reverse primer | GGGTGCACTGTATCGTGTGA | Minus | 20 | 4314 | 4295 | 60.04 | 55.00 | 6.00 | 1.00 |
| Product length | 141 | | | | | | | | |

**2 Gene name: XM_030474292.1 (NT-support)**

|  | Sequence (5'->3') | Tem. strand | Len | Start | End | Tm | GC% | Self com. | Self 3' com. |
| --- | --- | --- | --- | --- | --- | --- | --- | --- | --- |
| Forward primer | CTCTGGACGGGGAACATCTG | Plus | 20 | 703 | 722 | 59.82 | 60.00 | 2.00 | 2.00 |
| Reverse primer | GACCCTACATCTGGTTGGCG | Minus | 20 | 970 | 951 | 60.46 | 60.00 | 3.00 | 2.00 |
| Product length | 268 | | | | | | | | |

**3 Gene name: NM_204277.1 (NT-support)**

|  | Sequence (5'->3') | Tem. strand | Len | Start | End | Tm | GC% | Self com. | Self 3' com. |
| --- | --- | --- | --- | --- | --- | --- | --- | --- | --- |
| Forward primer | GTCCAGCAAGCCTGTGAGAT | Plus | 20 | 1069 | 1088 | 60.04 | 55.00 | 4.00 | 2.00 |
| Reverse primer | CCCGTCATCCTGCTTGAACT | Minus | 20 | 1284 | 1265 | 60.04 | 55.00 | 3.00 | 1.00 |
| Product length | 216 | | | | | | | | |

**4 Gene name: XM_037371960.1 (NT-support)**

|  | Sequence (5'->3') | Tem. strand | Len | Start | End | Tm | GC% | Self com. | Self 3' com. |
| --- | --- | --- | --- | --- | --- | --- | --- | --- | --- |
| Forward primer | CCTGGACGACATGCTCTACG | Plus | 20 | 886 | 905 | 60.25 | 60.00 | 4.00 | 2.00 |
| Reverse primer | CGGTGTCCTTGTAGAGGTGC | Minus | 20 | 1016 | 997 | 60.39 | 60.00 | 3.00 | 2.00 |
| Product length | 131 | | | | | | | | |

**5 Gene name: XR_006936251.1 (NT-support)**

|  | Sequence (5'->3') | Tem. strand | Len | Start | End | Tm | GC% | Self com. | Self 3' com. |
| --- | --- | --- | --- | --- | --- | --- | --- | --- | --- |
| Forward primer | AAGCAGATACTGCGTACCTATGA | Plus | 23 | 24688 | 24710 | 59.11 | 43.48 | 6.00 | 1.00 |
| Reverse primer | ACAGTGCTGCCGTATTCCAG | Minus | 20 | 24917 | 24898 | 60.39 | 55.00 | 4.00 | 2.00 |
| Product length | 230 | | | | | | | | |

**6 Gene name: XM_030970703.1 (NT-support)**

|  | Sequence (5'->3') | Tem. strand | Len | Start | End | Tm | GC% | Self com. | Self 3' com. |
| --- | --- | --- | --- | --- | --- | --- | --- | --- | --- |
| Forward primer | TGGACATCATGCGGGTGAAC | Plus | 20 | 177 | 196 | 60.68 | 55.00 | 5.00 | 2.00 |
| Reverse primer | CTTCCACCAGTACTTGCGCT | Minus | 20 | 322 | 303 | 60.32 | 55.00 | 6.00 | 2.00 |
| Product length | 146 | | | | | | | | |

**7 Gene name: CR391159.1 (NT-support)**

|  | Sequence (5'->3') | Tem. strand | Len | Start | End | Tm | GC% | Self com. | Self 3' com. |
| --- | --- | --- | --- | --- | --- | --- | --- | --- | --- |
| Forward primer | AAGCACTTGGTGATAGCGCA | Plus | 20 | 46 | 65 | 60.32 | 50.00 | 4.00 | 2.00 |
| Reverse primer | TCCTGTGCTAGTCCTACCCG | Minus | 20 | 303 | 284 | 60.40 | 60.00 | 4.00 | 2.00 |
| Product length | 258 | | | | | | | | |

**8 Gene name: NR_172493.1 (NT-support)**

|  | Sequence (5'->3') | Tem. strand | Len | Start | End | Tm | GC% | Self com. | Self 3' com. |
| --- | --- | --- | --- | --- | --- | --- | --- | --- | --- |
| Forward primer | GGAAGACCCCCAGTGTTGTC | Plus | 20 | 192 | 211 | 60.25 | 60.00 | 3.00 | 1.00 |
| Reverse primer | TTCAGTGTCAGCCATCGGTC | Minus | 20 | 310 | 291 | 60.04 | 55.00 | 3.00 | 1.00 |
| Product length | 119 | | | | | | | | |

**9 Gene name: XM_046942855.1 (NT-support)**

|  | Sequence (5'->3') | Tem. strand | Len | Start | End | Tm | GC% | Self com. | Self 3' com. |
| --- | --- | --- | --- | --- | --- | --- | --- | --- | --- |
| Forward primer | TTTCAGTGTGCGCTGTTTGG | Plus | 20 | 1397 | 1416 | 59.90 | 50.00 | 4.00 | 0.00 |
| Reverse primer | AGACAAGCAAACCATGGGGA | Minus | 20 | 1581 | 1562 | 59.52 | 50.00 | 6.00 | 0.00 |
| Product length | 185 | | | | | | | | |

**10 Gene name: XR_002111465.1 (NT-support)**

|  | Sequence (5'->3') | Tem. strand | Len | Start | End | Tm | GC% | Self com. | Self 3' com. |
| --- | --- | --- | --- | --- | --- | --- | --- | --- | --- |
| Forward primer | TTTTGCCCCTTCCATAGCGT | Plus | 20 | 9070 | 9089 | 59.96 | 50.00 | 2.00 | 0.00 |
| Reverse primer | AGGAGTTTAAGCACGACGCA | Minus | 20 | 9160 | 9141 | 59.97 | 50.00 | 4.00 | 0.00 |
| Product length | 91 | | | | | | | | |

**11 Gene name: NM_001389564.2 (NT-support)**

|  | Sequence (5'->3') | Tem. strand | Len | Start | End | Tm | GC% | Self com. | Self 3' com. |
| --- | --- | --- | --- | --- | --- | --- | --- | --- | --- |
| Forward primer | CATCCCCTGGACACCGTCAA | Plus | 20 | 144 | 163 | 61.84 | 60.00 | 3.00 | 3.00 |
| Reverse primer | ATCTCCATGCCTCAACTGGC | Minus | 20 | 356 | 337 | 60.11 | 55.00 | 4.00 | 2.00 |
| Product length | 213 | | | | | | | | |

**12 Gene name: XR_001470382.4 (NT-support)**

|  | Sequence (5'->3') | Tem. strand | Len | Start | End | Tm | GC% | Self com. | Self 3' com. |
| --- | --- | --- | --- | --- | --- | --- | --- | --- | --- |
| Forward primer | AGAAGAGGCTGTCTCCCACC | Plus | 20 | 755 | 774 | 60.91 | 60.00 | 5.00 | 0.00 |
| Reverse primer | ACTCACATAAAGGAGGGGGC | Minus | 20 | 970 | 951 | 59.08 | 55.00 | 3.00 | 2.00 |
| Product length | 216 | | | | | | | | |

**13 Gene name: NM_205119.1 (NT-support)**

|  | Sequence (5'->3') | Tem. strand | Len | Start | End | Tm | GC% | Self com. | Self 3' com. |
| --- | --- | --- | --- | --- | --- | --- | --- | --- | --- |
| Forward primer | GCGAGGGAAATCCTCGACTC | Plus | 20 | 114 | 133 | 60.25 | 60.00 | 6.00 | 3.00 |
| Reverse primer | TGTTGATGTGCTCCACTGCT | Minus | 20 | 303 | 284 | 59.89 | 50.00 | 3.00 | 0.00 |
| Product length | 190 | | | | | | | | |

**14 Gene name: XM_046930159.1 (NT-support)**

|  | Sequence (5'->3') | Tem. strand | Len | Start | End | Tm | GC% | Self com. | Self 3' com. |
| --- | --- | --- | --- | --- | --- | --- | --- | --- | --- |
| Forward primer | ATACCTGAAGGCGCACGAAA | Plus | 20 | 3746 | 3765 | 60.04 | 50.00 | 4.00 | 0.00 |
| Reverse primer | TGGTGTAAAGGACCGACTGC | Minus | 20 | 3851 | 3832 | 59.97 | 55.00 | 3.00 | 2.00 |
| Product length | 106 | | | | | | | | |

**15 Gene name: XM_046926203.1 (NT-support)**

|  | Sequence (5'->3') | Tem. strand | Len | Start | End | Tm | GC% | Self com. | Self 3' com. |
| --- | --- | --- | --- | --- | --- | --- | --- | --- | --- |
| Forward primer | CCCACTGGGCTGAGGTTG | Plus | 18 | 1398 | 1415 | 59.97 | 66.67 | 7.00 | 0.00 |
| Reverse primer | AGCTGGCTCAGATGCTGTTT | Minus | 20 | 1536 | 1517 | 59.96 | 50.00 | 4.00 | 0.00 |
| Product length | 139 | | | | | | | | |

**16 Gene name: XR_006935483.1 (NT-support)**

|  | Sequence (5'->3') | Tem. strand | Len | Start | End | Tm | GC% | Self com. | Self 3' com. |
| --- | --- | --- | --- | --- | --- | --- | --- | --- | --- |
| Forward primer | GGTCTTGCAGCTGTGTTTGG | Plus | 20 | 1114 | 1133 | 59.97 | 55.00 | 6.00 | 0.00 |
| Reverse primer | ACAGCATGAAGTCCAAGGGG | Minus | 20 | 1327 | 1308 | 59.96 | 55.00 | 4.00 | 1.00 |
| Product length | 214 | | | | | | | | |

**17 Gene name: BX929356.1 (NT-support)**

|  | Sequence (5'->3') | Tem. strand | Len | Start | End | Tm | GC% | Self com. | Self 3' com. |
| --- | --- | --- | --- | --- | --- | --- | --- | --- | --- |
| Forward primer | GGAGCTTTTCCCGAAGCGTT | Plus | 20 | 105 | 124 | 61.24 | 55.00 | 4.00 | 2.00 |
| Reverse primer | GGTTCAGGTGCCGATCCAC | Minus | 19 | 208 | 190 | 60.75 | 63.16 | 4.00 | 2.00 |
| Product length | 104 | | | | | | | | |

**18 Gene name: HG999715.3 (NT-support)**

|  | Sequence (5'->3') | Tem. strand | Len | Start | End | Tm | GC% | Self com. | Self 3' com. |
| --- | --- | --- | --- | --- | --- | --- | --- | --- | --- |
| Forward primer | GTGGGTCAGGGAGAACCTC | Plus | 19 | 403 | 421 | 59.02 | 63.16 | 3.00 | 3.00 |
| Reverse primer | ATTAACACGTTTCCGGTGTCC | Minus | 21 | 555 | 535 | 58.85 | 47.62 | 4.00 | 1.00 |
| Product length | 153 | | | | | | | | |

**19 Gene name: XM_046937669.1 (NT-support)**

|  | Sequence (5'->3') | Tem. strand | Len | Start | End | Tm | GC% | Self com. | Self 3' com. |
| --- | --- | --- | --- | --- | --- | --- | --- | --- | --- |
| Forward primer | ATGCATCGCTTCCAAAAGGT | Plus | 20 | 1943 | 1962 | 58.45 | 45.00 | 6.00 | 0.00 |
| Reverse primer | ACTCCCGGTGCTTTGAGC | Minus | 18 | 2153 | 2136 | 59.97 | 61.11 | 4.00 | 3.00 |
| Product length | 211 | | | | | | | | |

**20 Gene name: XM_419678.8 (NT-support)**

|  | Sequence (5'->3') | Tem. strand | Len | Start | End | Tm | GC% | Self com. | Self 3' com. |
| --- | --- | --- | --- | --- | --- | --- | --- | --- | --- |
| Forward primer | GGCTCGTTCAGATGCAAAGT | Plus | 20 | 68 | 87 | 58.84 | 50.00 | 4.00 | 1.00 |
| Reverse primer | CTGCTCTCTGGTGACAGGAA | Minus | 20 | 228 | 209 | 59.03 | 55.00 | 4.00 | 1.00 |
| Product length | 161 | | | | | | | | |

**21 Gene name: XM_040688341.2 (NT-support)**

|  | Sequence (5'->3') | Tem. strand | Len | Start | End | Tm | GC% | Self com. | Self 3' com. |
| --- | --- | --- | --- | --- | --- | --- | --- | --- | --- |
| Forward primer | AGAAGGGTGCATCATAGCAGT | Plus | 21 | 4805 | 4825 | 59.16 | 47.62 | 5.00 | 1.00 |
| Reverse primer | CTTGCATTTACCAGGCCACC | Minus | 20 | 5052 | 5033 | 59.47 | 55.00 | 4.00 | 1.00 |
| Product length | 248 | | | | | | | | |

**22 Gene name: XR_005843481.2 (NT-support)**

|  | Sequence (5'->3') | Tem. strand | Len | Start | End | Tm | GC% | Self com. | Self 3' com. |
| --- | --- | --- | --- | --- | --- | --- | --- | --- | --- |
| Forward primer | GGTTCGGATTGCCTCTTTGC | Plus | 20 | 2791 | 2810 | 59.83 | 55.00 | 3.00 | 2.00 |
| Reverse primer | GCAGAGCAATGCACAAGACC | Minus | 20 | 3037 | 3018 | 60.11 | 55.00 | 5.00 | 0.00 |
| Product length | 247 | | | | | | | | |

**23 Gene name: XM_005533218.2 (NT-support)**

|  | Sequence (5'->3') | Tem. strand | Len | Start | End | Tm | GC% | Self com. | Self 3' com. |
| --- | --- | --- | --- | --- | --- | --- | --- | --- | --- |
| Forward primer | GTCCGGAAAGTGTCCGATGT | Plus | 20 | 383 | 402 | 60.04 | 55.00 | 7.00 | 0.00 |
| Reverse primer | CGTACCCGTGATGAAGCTGA | Minus | 20 | 481 | 462 | 59.83 | 55.00 | 4.00 | 1.00 |
| Product length | 99 | | | | | | | | |

**24 Gene name: XM_040539289.1 (NT-support)**

|  | Sequence (5'->3') | Tem. strand | Len | Start | End | Tm | GC% | Self com. | Self 3' com. |
| --- | --- | --- | --- | --- | --- | --- | --- | --- | --- |
| Forward primer | CGTTCCAGAACTACCCCGAC | Plus | 20 | 890 | 909 | 60.11 | 60.00 | 6.00 | 1.00 |
| Reverse primer | TGGAAGAACGTGTCCACCTG | Minus | 20 | 968 | 949 | 59.89 | 55.00 | 4.00 | 2.00 |
| Product length | 79 | | | | | | | | |

**25 Gene name: AC186848.2 (NT-support)**

|  | Sequence (5'->3') | Tem. strand | Len | Start | End | Tm | GC% | Self com. | Self 3' com. |
| --- | --- | --- | --- | --- | --- | --- | --- | --- | --- |
| Forward primer | TGCACTGACCTTGGTGGTTT | Plus | 20 | 354 | 373 | 60.03 | 50.00 | 4.00 | 0.00 |
| Reverse primer | TCAGGTAAGAGCCAGCTCCA | Minus | 20 | 543 | 524 | 60.25 | 55.00 | 6.00 | 2.00 |
| Product length | 190 | | | | | | | | |

**26 Gene name: XM_031607112.1 (NT-support)**

|  | Sequence (5'->3') | Tem. strand | Len | Start | End | Tm | GC% | Self com. | Self 3' com. |
| --- | --- | --- | --- | --- | --- | --- | --- | --- | --- |
| Forward primer | TGTGAGGTCCCAGTTTACGG | Plus | 20 | 53 | 72 | 59.32 | 55.00 | 4.00 | 2.00 |
| Reverse primer | CCACCTGACCGGAAGTGAAG | Minus | 20 | 237 | 218 | 60.32 | 60.00 | 4.00 | 0.00 |
| Product length | 185 | | | | | | | | |

**27 Gene name: XM_014262261.1 (NT-support)**

|  | Sequence (5'->3') | Tem. strand | Len | Start | End | Tm | GC% | Self com. | Self 3' com. |
| --- | --- | --- | --- | --- | --- | --- | --- | --- | --- |
| Forward primer | CATGATCGTCACCGAGTTCCT | Plus | 21 | 1849 | 1869 | 59.86 | 52.38 | 4.00 | 1.00 |
| Reverse primer | ACACTTTGCAGACCAGGTGA | Minus | 20 | 2051 | 2032 | 59.45 | 50.00 | 5.00 | 1.00 |
| Product length | 203 | | | | | | | | |

**28 Gene name: XM_040652639.2 (NT-support)**

|  | Sequence (5'->3') | Tem. strand | Len | Start | End | Tm | GC% | Self com. | Self 3' com. |
| --- | --- | --- | --- | --- | --- | --- | --- | --- | --- |
| Forward primer | GAAGCGACGGCAAGATCCA | Plus | 19 | 2480 | 2498 | 60.45 | 57.89 | 4.00 | 0.00 |
| Reverse primer | CTATCAGCGGCGTGAGAAGGG | Minus | 21 | 2560 | 2540 | 62.87 | 61.90 | 5.00 | 0.00 |
| Product length | 81 | | | | | | | | |

**29 Gene name: XM_046906388.1 (NT-support)**

|  | Sequence (5'->3') | Tem. strand | Len | Start | End | Tm | GC% | Self com. | Self 3' com. |
| --- | --- | --- | --- | --- | --- | --- | --- | --- | --- |
| Forward primer | CCAGGCACACCAGTTCAAAG | Plus | 20 | 74 | 93 | 59.33 | 55.00 | 3.00 | 0.00 |
| Reverse primer | AGGATCACAACCCCAAAGCC | Minus | 20 | 352 | 333 | 60.25 | 55.00 | 4.00 | 1.00 |
| Product length | 279 | | | | | | | | |

**30 Gene name: XM_040695062.2 (NT-support)**

|  | Sequence (5'->3') | Tem. strand | Len | Start | End | Tm | GC% | Self com. | Self 3' com. |
| --- | --- | --- | --- | --- | --- | --- | --- | --- | --- |
| Forward primer | GCTTGGAGGTGCTCTTCAGT | Plus | 20 | 4808 | 4827 | 59.96 | 55.00 | 4.00 | 1.00 |
| Reverse primer | CGGCAGAATAACCTCCTTTGC | Minus | 21 | 5037 | 5017 | 59.60 | 52.38 | 3.00 | 2.00 |
| Product length | 230 | | | | | | | | |

**31 Gene name: XM_046937999.1 (NT-support)**

|  | Sequence (5'->3') | Tem. strand | Len | Start | End | Tm | GC% | Self com. | Self 3' com. |
| --- | --- | --- | --- | --- | --- | --- | --- | --- | --- |
| Forward primer | TCTGGCTGGGAAATAGGCTC | Plus | 20 | 7200 | 7219 | 59.16 | 55.00 | 3.00 | 1.00 |
| Reverse primer | GCAATCCAGAAGCCATCCAA | Minus | 20 | 7304 | 7285 | 58.52 | 50.00 | 3.00 | 0.00 |
| Product length | 105 | | | | | | | | |

**32 Gene name: XM_046915922.1 (NT-support)**

|  | Sequence (5'->3') | Tem. strand | Len | Start | End | Tm | GC% | Self com. | Self 3' com. |
| --- | --- | --- | --- | --- | --- | --- | --- | --- | --- |
| Forward primer | ATTGCCGATTCCTCCACAGG | Plus | 20 | 1228 | 1247 | 60.11 | 55.00 | 3.00 | 2.00 |
| Reverse primer | CGTGCACTCCAAACACCAAG | Minus | 20 | 1410 | 1391 | 59.97 | 55.00 | 6.00 | 0.00 |
| Product length | 183 | | | | | | | | |

**33 Gene name: Gal6 (Novel)**

|  | Sequence (5'->3') | Tem. strand | Len | Start | End | Tm | GC% | Self com. | Self 3' com. |
| --- | --- | --- | --- | --- | --- | --- | --- | --- | --- |
| Forward primer | GGACCGGAAGCGGAAGTAGA | Plus | 20 | 151 | 170 | 61.32 | 60.00 | 4.00 | 0.00 |
| Reverse primer | AATGGAGCGGTTTCGGGTTG | Minus | 20 | 284 | 265 | 61.24 | 55.00 | 2.00 | 0.00 |
| Product length | 134 | | | | | | | | |

**34 Gene name: Gal1779 (Novel)**

|  | Sequence (5'->3') | Tem. strand | Len | Start | End | Tm | GC% | Self com. | Self 3' com. |
| --- | --- | --- | --- | --- | --- | --- | --- | --- | --- |
| Forward primer | TAACGACCCCCACGGTCAC | Plus | 19 | 20 | 38 | 61.57 | 63.16 | 4.00 | 3.00 |
| Reverse primer | ATGGAGATGGCCGTGTTTGC | Minus | 20 | 149 | 130 | 61.32 | 55.00 | 4.00 | 2.00 |
| Product length | 130 | | | | | | | | |

**35 Gene name: Gal2189 (Novel)**

|  | Sequence (5'->3') | Tem. strand | Len | Start | End | Tm | GC% | Self com. | Self 3' com. |
| --- | --- | --- | --- | --- | --- | --- | --- | --- | --- |
| Forward primer | TTTGGGGTCTTCACGGTGTT | Plus | 20 | 69 | 88 | 59.74 | 50.00 | 5.00 | 1.00 |
| Reverse primer | CTCACTTTGGGTCCTGGCTT | Minus | 20 | 231 | 212 | 59.89 | 55.00 | 3.00 | 0.00 |
| Product length | 163 | | | | | | | | |

**36 Gene name: Gal559 (Novel)**

|  | Sequence (5'->3') | Tem. strand | Len | Start | End | Tm | GC% | Self com. | Self 3' com. |
| --- | --- | --- | --- | --- | --- | --- | --- | --- | --- |
| Forward primer | CCCTGGAGAATGGCCGAATG | Plus | 20 | 160 | 179 | 60.82 | 60.00 | 4.00 | 0.00 |
| Reverse primer | CCAAATGCAGCCAGAACTCG | Minus | 20 | 343 | 324 | 59.83 | 55.00 | 4.00 | 2.00 |
| Product length | 184 | | | | | | | | |

**37 Gene name: Gal2405 (Novel)**

|  | Sequence (5'->3') | Tem. strand | Len | Start | End | Tm | GC% | Self com. | Self 3' com. |
| --- | --- | --- | --- | --- | --- | --- | --- | --- | --- |
| Forward primer | CCATAAAGGGGGCGATTTCCA | Plus | 21 | 152 | 172 | 60.41 | 52.38 | 3.00 | 0.00 |
| Reverse primer | GAACTACTGACGCTACCCCA | Minus | 20 | 275 | 256 | 58.82 | 55.00 | 2.00 | 0.00 |
| Product length | 124 | | | | | | | | |

**38 Gene name: Gal81 (Novel)**

|  | Sequence (5'->3') | Tem. strand | Len | Start | End | Tm | GC% | Self com. | Self 3' com. |
| --- | --- | --- | --- | --- | --- | --- | --- | --- | --- |
| Forward primer | TACCGCTTTCAGGCAGTAGC | Plus | 20 | 94 | 113 | 60.11 | 55.00 | 6.00 | 2.00 |
| Reverse primer | GTACCCCCAAGCGATACAGG | Minus | 20 | 208 | 189 | 59.89 | 60.00 | 4.00 | 0.00 |
| Product length | 115 | | | | | | | | |

**39 Gene name: Gal12 (Novel)**

|  | Sequence (5'->3') | Tem. strand | Len | Start | End | Tm | GC% | Self com. | Self 3' com. |
| --- | --- | --- | --- | --- | --- | --- | --- | --- | --- |
| Forward primer | GGTCGGCTCAGATGGGTTC | Plus | 19 | 155 | 173 | 60.15 | 63.16 | 3.00 | 0.00 |
| Reverse primer | CAATCTGAACTGGGCTGAGGT | Minus | 21 | 229 | 209 | 60.00 | 52.38 | 3.00 | 2.00 |
| Product length | 75 | | | | | | | | |

**40 Gene name: Gal7 (Novel)**

|  | Sequence (5'->3') | Tem. strand | Len | Start | End | Tm | GC% | Self com. | Self 3' com. |
| --- | --- | --- | --- | --- | --- | --- | --- | --- | --- |
| Forward primer | AGGGGTTCGCTTTGGATTGG | Plus | 20 | 215 | 234 | 60.61 | 55.00 | 2.00 | 0.00 |
| Reverse primer | GGCTCCCATAGGTGGAGATG | Minus | 20 | 299 | 280 | 59.31 | 60.00 | 6.00 | 0.00 |
| Product length | 85 | | | | | | | | |

**41 Gene name: Gal385 (Novel)**

|  | Sequence (5'->3') | Tem. strand | Len | Start | End | Tm | GC% | Self com. | Self 3' com. |
| --- | --- | --- | --- | --- | --- | --- | --- | --- | --- |
| Forward primer | AAAGAGGGCAGAAAAGTGGGC | Plus | 21 | 26 | 46 | 61.10 | 52.38 | 2.00 | 2.00 |
| Reverse primer | GAAGCTTTCTGTGGCCCTTCT | Minus | 21 | 212 | 192 | 60.55 | 52.38 | 6.00 | 0.00 |
| Product length | 187 | | | | | | | | |

**42 Gene name: Gal556 (Novel)**

|  | Sequence (5'->3') | Tem. strand | Len | Start | End | Tm | GC% | Self com. | Self 3' com. |
| --- | --- | --- | --- | --- | --- | --- | --- | --- | --- |
| Forward primer | TTCTCCCGAAGAAATCGTGGC | Plus | 21 | 8 | 28 | 60.67 | 52.38 | 7.00 | 2.00 |
| Reverse primer | CTCCTCTTCGTGCTCTCTGTT | Minus | 21 | 279 | 259 | 59.46 | 52.38 | 2.00 | 0.00 |
| Product length | 272 | | | | | | | | |

**actin-β:**

| F:gtgtgatggttggtatgggc | Length: 20 bp |
| --- | --- |
| R:ctctgttggctttggggttc | Length: 20 bp |
